# SARS-CoV-2 infection of circulating immune cells is not responsible for virus dissemination in severe COVID-19 patients

**DOI:** 10.1101/2021.01.19.427282

**Authors:** Nicole L. Rosin, Arzina Jaffer, Sarthak Sinha, Rory P. Mulloy, Carolyn Robinson, Elodie Labit, Luiz G. Almeida, Antoine Dufour, Jennifer A. Corcoran, Bryan Yipp, Jeff Biernaskie

**Author notes:** Corresponding authors Jeff Biernaskie –, Bryan Yipp –, Nicole Rosin –.

## Abstract

In late 2019 a novel coronavirus (SARS-CoV-2) emerged, and has since caused a global pandemic. Understanding the pathogenesis of COVID-19 disease is necessary to inform development of therapeutics, and management of infected patients. Using scRNAseq of blood drawn from SARS-CoV-2 patients, we asked whether SARS-CoV-2 may exploit immune cells as a ‘Trojan Horse’ to disseminate and access multiple organ systems. Our data suggests that circulating cells are not actively infected with SARS-CoV-2, and do not appear to be a source of viral dissemination.

## Introduction

Coronavirus 2019 (COVID-19), the disease caused by Severe Acute Respiratory Syndrome Coronavirus 2 (SARS-CoV-2) was first identified in late 2019, but by December 9, 2020 has spread to 220 countries, with over 67 million confirmed cases and over 1.5 million confirmed deaths ^1^. Patients with COVID-19 present with a diverse range of symptoms from asymptomatic, mild (cough, fatigue), moderate (non-mild pneumonia), severe (dyspnea, low oxygen saturation) to critical (respiratory failure, septic shock and/or multiple organ dysfunction or failure) ^2,3^. In addition to this range of respiratory symptoms, there are emerging reports of thromboembolism ^4^, neural defects (both central and peripheral, ie. anosmia and ageusia ^5,6^), and Kawasaki-like symptoms in children ^7^. Currently, the pathophysiology of COVID-19, and more specifically how the immune response to SARS-CoV-2 contributes to clinical disease progression, remains poorly understood.

Coronaviruses are enveloped, non-segmented, positive-sense RNA viruses that can infect both humans and many other mammals ^3^. Of those that infect humans, some cause only mild symptoms, such as hCoV-OC43 ^8^. This ‘common cold’ coronavirus shares only 56% genomic sequence homology with SARS-CoV-2 (using BLAST search: https://blast.ncbi.nlm.nih.gov/Blast.cgi). However, other coronaviruses cause severe respiratory diseases; Severe acute respiratory syndrome coronavirus (SARS-CoV) was discovered in 2003 and causes SARS, while Middle East respiratory syndrome coronavirus (MERS-CoV) was discovered in 2012 and causes MERS. SARS-CoV and MERS-CoV infection have 10% and 37% mortality rates respectively ^9,10^ and share 79% and 50% genomic sequence homology to SARS-CoV-2, respectively ^11^.

SARS-CoV-2 has been identified in the blood of infected patients ^12^, as have SARS-CoV ^13,14^ and MERS-CoV ^15,16^. Conflicting reports exist regarding viremia (infectious virus in the circulation) in COVID-19, ^17,18^ versus the presence of viral RNA in blood samples ^3^ and the clinical significance of this nuance. Indeed, previous reports of viremia in COVID-19 patients include those using bulk RNAseq ^19,20^, which fail to convincingly show that individual cells of the circulation contain SARS-CoV-2 genetic material.

A recent pre-print reported SARS-CoV-2 nucleoprotein in CD68+/CD169+ macrophages in the lungs of severe COVID-19 patients ^21^, supporting a previous report of SARS-CoV-2 RNA containing macrophages in the bronchoalveolar lavage fluid (BALF) ^22^. Furthermore, multiple organ systems beyond the lung are also affected in severe COVID-19 disease ^23^, and multiorgan tropism has been reported ^24^, which raises the possibility that visceral organ infection may be contributing to disease severity, though a causal link has not yet been demonstrated. In light of this, a review by *Abassi* and colleagues recently questioned whether SARS-CoV-2 dissemination to distal organs is facilitated by infected macrophages re-entering the circulation ^25^. Here, our objective was to understand the potential for circulating immune cell infection as a mode of viral dissemination to distal organs.

## Results

### Viral Protein Circulates in Severe COVID-19 Patients

Five patients with severe acute respiratory distress syndrome (ARDS) and PCR confirmed SARS-CoV-2 infection (severe COVID-19 disease) were enrolled and had blood drawn within 72 h of admission to the ICU (Supplementary Table 1). We detected one or more viral proteins in the plasma of each of the five patients (Table 1). NSP3 (encoded by the ORF1ab gene) was detected in all samples, while NSP3, the spike protein S1 and RNA-directed RNA polymerase were found in three of the plasma samples. One sample, from patient UC_5, contained 5 SARS-CoV-2 specific proteins.

**Table 1.** Proteomics of SARS-CoV-2 in patient plasma

| Sequence | SARS-CoV-2 Protein | Reporter Ion Intensity |  |  |  |  |
| --- | --- | --- | --- | --- | --- | --- |
|  |  | UC_4 | UC_5 | UC_6 | UC_7 | UC_8 |
| EFVFKNIDGYFK | Spike protein S1 | 0 | 20977 | 12262 | 21872 | 0 |
| SYELQTPFEIKLAK | non-structural protein 2 | 0 | 34243 | 0 | 0 | 0 |
| AFKQIVESC GNFK | non-structural protein 2 | 0 | 5898.9 | 0 | 0 | 0 |
| VTFGDDTVIEVQGYK | non-structural protein 2 | 0 | 15652 | 0 | 0 | 0 |
| NLYDKLVSSFLEMK | non-structural protein 3 | 23313 | 56518 | 44610 | 73525 | 18845 |
| GFFKEGSSVELK | RNA-directed RNA polymerase | 0 | 8380.8 | 0 | 0 | 0 |
| SVLYYQNNVFMSEAK | RNA-directed RNA polymerase | 0 | 26229 | 21261 | 19505 | 0 |
| NVATLQAENV TGLFK | proofreading exoribonuclease | 0 | 16095 | 0 | 0 | 0 |
| PPPGDQFKHLIPLMYK | proofreading exoribonuclease | 0 | 13779 | 0 | 0 | 0 |

### SARS-CoV-2 Does Not Infect Circulating Immune Cells

From whole blood, we isolated leukocytes and enriched for the lymphocyte population to ensure that enough lymphocytes would be collected/assessed due to reported lymphocytopenia ^26^. The leukocytes and lymphocytes were mixed in equal ratios prior to conducting scRNAseq using the 10X Genomics Single Cell 3’ NextGEM (v3.1) technology. The resulting sequences were aligned with the human genome (GRCh38) appended with the SARS-CoV-2 genome. Two versions of the SARS-CoV-2 reference genome were used to account for potential shifting in the sequence over time and possibility of the predominant variant differing based on geographical locations. MT412228 was isolated in Seattle, USA and MN908947.3 was isolated in Shenzhen, China. Because our study was conducted in Calgary, Alberta, Canada, MT412228 is reported here unless specified, as at the time of publication this reference genome was the closest available geographically to Calgary. This method of SARS-CoV-2 RNA detection in the peripheral blood cell samples was confirmed by aligning a control group of publicly available bronchoalveolar lavage fluid (denoted by BALF_) sample sequences to the same custom reference genome. These BALF samples were isolated from three healthy control patients, three mild COVID-19 patients, and six patients with severe COVID-19 from Shenzhen, China ^27^.

All UC_ and BALF_ samples were aggregated together using CellRanger, and processed using the Seurat workflow ^28^. Supplementary Table 2 outlines the number of cells aggregated from each sample after quality control. The peripheral immune cells clustered together, but did not overlap with the BALF samples, as expected. Distinct cell types were annotated based on established cell type markers, with UC_ denoting cell types found only within the peripheral immune cell samples (Figure 1).

**Figure 1.**
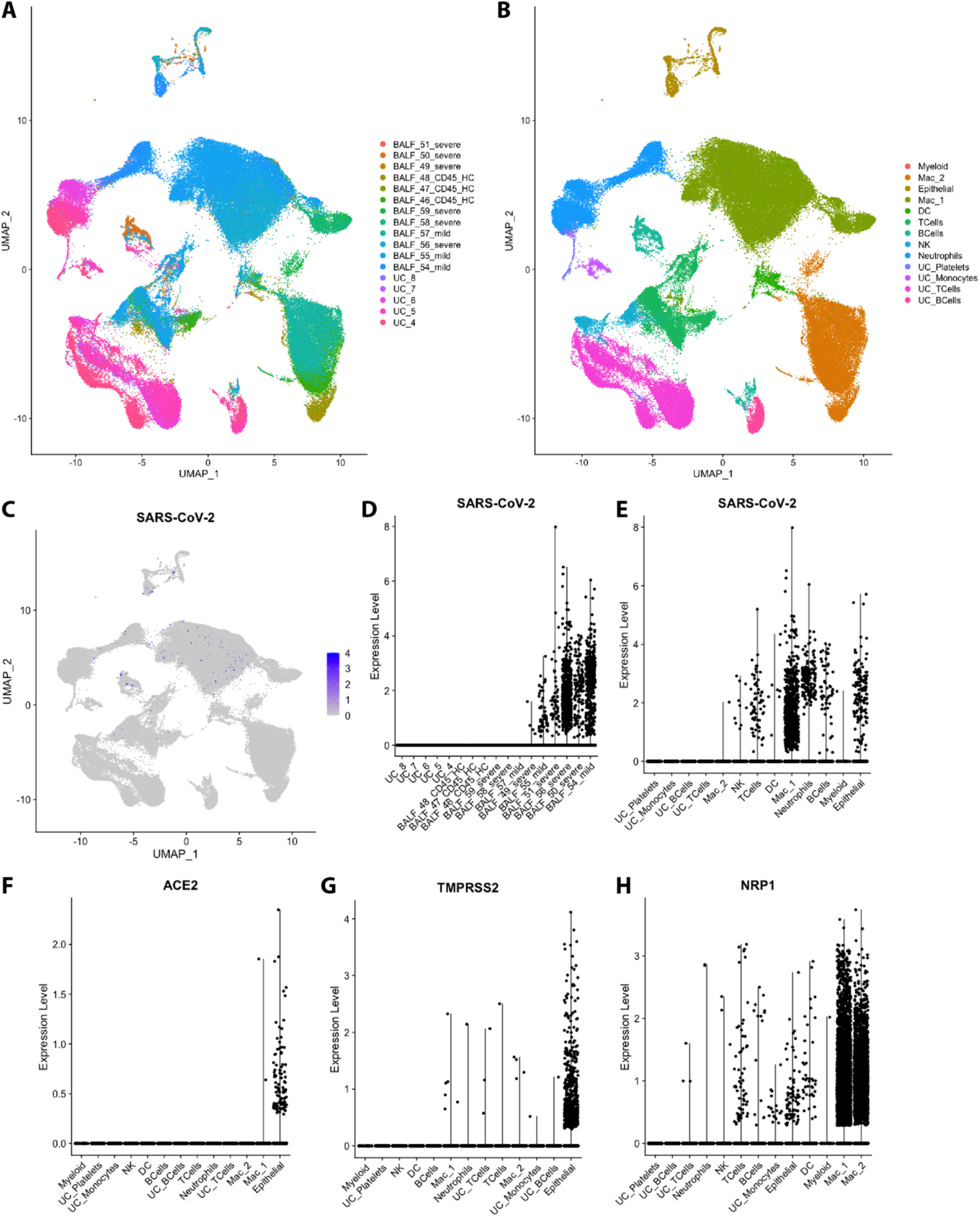
SARS-CoV-2 RNA (MT412228) is detected in BALF but not circulating immune cells. A) UMAP of all sample sequences coloured by sample ID (BALF or UC denoting origin of sample) B) UMAP of all sample sequences coloured by sample type C) UMAP illustrating the detection of SARS-CoV-2 RNA D and E) Violin plots illustrating detection of SARS-CoV-2 RNA according to sample ID and cell type F, G and H) Violin plots denoting expression of ACE2, TMPRSS2 and NRP1 expression according to cell type

We found that none of the five mixed leukocyte/lymphocyte samples (denoted by UC_) contained any SARS-CoV-2 RNA (Figure 1, Supplementary Table 3). However, as discussed by *Bost et al*, the BALF samples contained SARS-CoV-2 RNA, which was located primarily in macrophages and epithelial cells ^22^. Using our custom genome as a reference, six of the nine COVID-19 BALF samples (two mild and four severe) contain SARS-CoV-2 RNA (Figure 1, Supplementary Table 3), again in concordance with the *Bost et al* study. The same samples were positive when aligned to the control reference genome containing MN908947.3 (Figure 2). This suggests that aligning to our custom SARS-CoV-2 reference genome is a robust method of detecting SARS-CoV-2, and taken together supports that peripheral immune cells do not contain SARS-CoV-2 RNA. Of note, we examined one additional SARS-CoV-2 positive patient who presented with a thrombotic stroke, but no respiratory symptoms or ARDS (and subsequently was not included in the aggregated data presented here), who also lacked any detectable SARS-CoV-2 RNA in the isolated circulating immune cells. While anecdotal, given we studied only one patient exhibiting clotting symptoms, it does suggest that this divergent clinical presentation is also not due to direct infection of circulating platelets or other immune cells.

**Figure 2.**
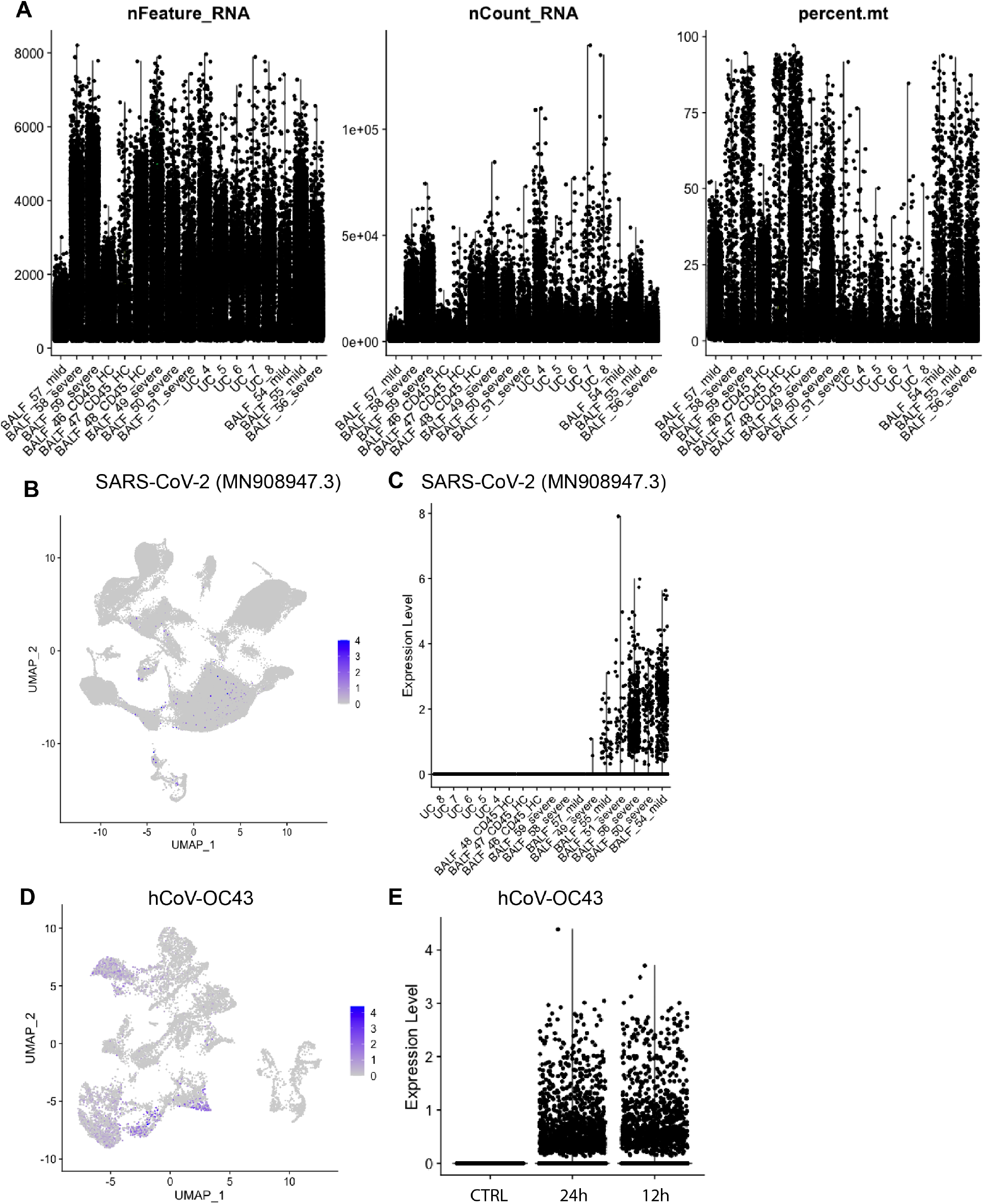
Quality control metrics, SARS-CoV-2 (MN908947.3) and hCoV-OC43 detection. A) Quality Control Metrics for all samples aligned to MT412228, including nFeature (number of genes), nCount (number of UMI), precent.mt (percent of mitochrondrial RNA). All of the samples were aligned to a custom reference genome consisting of GRCh38-3.0.0 with MN908947.3 appended B) UMAP illustrating the detection of SARS-CoV-2 RNA (aligned to MN908947.3), and C) Violin plot illustrating detection of SARS-CoV-2 RNA (aligned to MN908947.3) according to sample ID. A second dataset was generated from HUVECs infected with hCoV-OC43 at a titre of 3.5×10^4^ TCID50/mL for either 12h or 24h. D) UMAP illustrating the detection of hCoV-OC43 RNA in aggregated samples, and E) Violin plot illustrating detection of hCoV-OC43 RNA according to length of infection.

### Immune Cells in the Lung Contain SARS-CoV-2 RNA

SARS-CoV-2 was detected in cell types harboring surface receptors that facilitate SARS-CoV-2 entry into human cells. Epithelial cells, macrophages and neutrophils were the three most common cell types in which SARS-CoV-2 was detected (Figure 1, Supplementary Table 4). Unsurprisingly, the spike protein cell surface processing protease transmembrane protease serine 2 (TMPRSS2) and surface receptor (for host cell entry) angiotensin-converting enzyme 2 (ACE2) ^29^ were highly expressed in epithelial cells (Figure 1). Macrophages expressed neuropilin-1 (NRP1), which has also been reported as a means of host cell entry (Figure 1)^30,31^. Neutrophils do not express ACE2, TMPRSS2 or NRP1, suggesting that they phagocytosed viral particles or their products without being actively infected, and raises the possibility that macrophages may do the same.

It is also important to note that the BALF samples were isolated using 10X Genomics 5’ technology, whereas the peripheral immune cells presented here, were isolated using 10X Genomics 3’ technology. Identification of SARS-CoV-2 RNA should theoretically be captured using either technology. However, to ensure that coronavirus RNA could be detected in the peripheral cells, we also prepared a control sample of hCoV-OC43 (a ‘common cold’ coronavirus)-infected primary Human Umbilical Vein Endothelial Cells (HUVECs, Lonza) as a control for 3’ capture of viral RNA. OC43-infected HUVECs were processed identically to the peripheral cells. We aligned the resulting sequences with a custom reference genome, again based on GRCh38, with the hCoV-OC43 genome (NC_006213.1) appended, and were able to reliably detect hCoV-OC43 RNA using our analysis pipeline (Figure 2), suggesting that the inability to detect SARS-CoV-2 in the circulating immune cells was not due to the 3’ sequencing approach.

## Discussion

We identified viral protein in the plasma of all five of the patients in this study, which raises the question of whether the viral load is correlated with the patient’s clinical outcome. As mentioned, we detected five viral proteins in the plasma from patient UC_5, more than in the other four patients. Patient UC_5 was the only participant enrolled in our study that died while in the ICU. This recapitulates a recently published association between viral RNA load and COVID-19 severity and mortality, in which approximately a third of plasma samples contained detectable SARS-CoV-2 RNA^32^. While our sample size is limited, the fact that we identified viral proteins in the plasma from all five participants suggests that either, our participants had more severe disease and/or the LC-MS/MS method used here may be a more sensitive detection method for SARS-CoV-2 in plasma. In either case, this warrants further investigation.

Our study was not designed to determine if there is a causal link between viral tropism and disease severity or symptoms in distal organs, and there is still no direct evidence of such in literature. Causality of a direct viral effect in distal organs is difficult to parse out, partly due to the systemic effects of the immune response to severe lung infection. However, this remains an important concern that could directly affect treatment options for patients with severe COVID-19.

While immune cells in the lung are likely infected based on previous reports, our data suggests that the circulation of free virus, but not SARS-CoV-2 infected circulating immune cells, is the primary route of viral dissemination to visceral organs.

## Methods

### Enrolment and consent/REB approval

Peripheral blood samples were obtained from consenting ICU patients, recovered COVID-19, and healthy donors in Calgary, Canada. All experiments involving or human samples received approval from the Conjoint Health Ethics Review Board at the University of Calgary.

### Lymphocyte preparation

Whole blood (2mL) was collected into 5mL polystyrene round-bottom tubes. Tubes were spun (15min, 3000rpm, Room Temperature [RT]), plasma removed, and stored at-80°C for plasma proteomic analysis. 100*μ*L of Isolation Cocktail and 100*μ*L of Rapid Spheres (Easy Sep™ Direct Human Total Lymphocytes Isolation Kit: 19655, StemCell Technologies) were added to the remaining whole blood. After mixing and 5min incubation at RT, the sample volumes of ≤ 2.5mL blood were topped up to 5mL with D-PBS+2%FBS + 1mM EDTA. The diluted sample was incubated in the magnet without lid for 5min, at RT. This last step was repeated twice before cell resuspension in 5mL of PBS+0.04% BSA. After 2 washes in PBS+0.04% BSA, and centrifugation for 5 min at 2000rpm, 7500 lymphocytes were resuspended in 25*μ*L of PBS+0.04% BSA.

### Leukocyte preparation

Whole blood was collected in heparin containing vacuutubes and 12*μ*L of 0.5M EDTA with 1mL of PBS+2% FBS and 50*μ*L of RBC of EasySep RBC Depletion spheres (EasySep ™ RBC Depletion Reagent: 18170, Stem Cell Technologies) were added. After 5 min of magnet incubation, at RT, tubes were inverted and poured into a new tube. 50*μ*L of RBC depletion spheres were added. 5 min after incubation on the magnet, the tube was inverted and cell suspension was poured into a new 15mL tube. After 2 washes in 5mL of PBS+0.04% BSA and centrifugation at 2000rpm for 5 min at 20°C, cells were resuspended in 25*μ*L of PBS+0.04% BSA. A mixture of 7500 total leukocytes and 7500 lymphocytes were loaded for each patient into the 10X chip for scRNAseq.

### hCoV-OC43 preparation and HUVEC transduction

Stocks of hCoV-OC43 (ATCC) were propagated in Vero E6 cells (ATCC). To produce viral stocks, Vero E6 cells were infected at an MOI of 0.01 for 1h in serum-free DMEM (Thermo Fisher) at 33°C in a humidified incubator with 5% CO_2_. Following infection, the viral inoculum was removed and replaced with DMEM supplemented with 2% heat-inactivated FBS (Thermo Fisher) and 100 units/mL penicillin/streptomycin/glutamine (Thermo Fisher). After 6 days, the supernatant was harvested and centrifuged at 2000 RPM for 5 mins to isolate cell-free virus. Virus stocks were stored at-80°C. Viral titres were enumerated using Reed and Muench tissue-culture infectious dose 50% (TCID50) in Vero E6 cells ^33^.

Pooled human umbilical cord vein endothelial cells (HUVECs, Lonza, Basal, Switzerland) were cultured using endothelial growth media (EGM-2; Lonza). To passage cells, cell culture plates were pre-coated with 0.1% (w/v) porcine gelatine (Sigma, St. Louis, Missouri, US) in 1X PBS for 30 min at 37°C in a humidified incubator with 5% CO_2_. For infection, HUVECs were seeded into a pre-coated plate to achieve ∼80% confluency after 24 hours. The following day, the growth media was removed and replaced with 100mL of hCoV-OC43 inoculum and incubated at 37°C for one hour, shaking the plate every 10 minutes to distribute viral inoculum. Following incubation, the virus inoculum was removed and replaced with fresh EGM-2 for 12 or 24 hours before cells were prepared for scRNAseq.

### Reference genome construction, alignment and aggregation

Raw sequencing reads (BCL files) were converted to FASTQs using the CellRanger mkfastq pipeline (version 3.0.1). Transcript alignment was performed against a modified human reference transcriptome generated using CellRanger’s mkref pipeline by appending either MT412228 (https://www.ncbi.nlm.nih.gov/nuccore/MT412228) or MN908947.3 (https://www.ncbi.nlm.nih.gov/nuccore/MN908947) sequence to a pre-built GRCh38-3.0.0 reference. SARS-CoV-2 GTF files were generated with the feature type marked as “exon” which were concatenated with GRCh38-3.0.0 GTF annotation files. Likewise, the SARS-CoV-2 FASTAs containing raw sequences were concatenated with the genome.fa file in GRCh38-3.0.0 reference. FASTA and GTF outputs following concatenation were provided as inputs to “fasta” and “genes” arguments for cellranger mkref. Note that this approach did not generate a conjoined genome containing both human and SARS-CoV-2 sequences as separate genomes. Instead this approach added a new pseudogene called “SARS-CoV-2” to the human genome, as SARS-CoV-2 is expected to generate transcripts in infected human cells. Reads from individual libraries were processed using Cellranger count and were then concatenated into a single feature-barcode matrix using Cellranger aggr, with “mapped” normalization mode to subsample reads from higher-depth GEMs to ensure each library received equivalent read count (Supplementary Table 2).

### Publicly available control samples and re-alignment

Raw single-cell BALF samples isolated in Shenzhen, China, as described in *Liao, Liu, Yuan, et al*. and accessioned under SRP250732 were downloaded using the SRA-Toolkit’s prefetch function ^27^. The following samples were downloaded: SRR11181959 (severe COVID), SRR11181956 (severe COVID), SRR11181958 (severe COVID), SRR11537947 (healthy control), SRR11181954 (mild COVID), SRR11181955 (mild COVID), SRR11537946 (healthy control), SRR11181957 (mild COVID), SRR11537948 (healthy control), SRR11537949 (severe COVID), SRR11537951 (severe COVID), and SRR11537950 (severe COVID). SRA files were converted to FASTQs using the fastq-dump--split-files function in SRA Toolkit 2.10.5. FASTQs were aligned and aggregated with peripheral blood scRNA-Seq samples by mapping to our custom (SARS-CoV-2-appended to GRCh38-3.0.0) assembly, as described above.

### Dimensionality reduction, clustering and gene expression

Output aggregated, filtered feature matrix files were imported into a SeuratObject using Seurat V3.1.5 and R Version 3.6.1, using default parameters ^28,34^. Data was subsetted on ‘nCount_RNA’, ‘nFeature_RNA’, and ‘percent.mt’ values of 30,000, 5,000 and 10% respectively. Normalization and scaling were done using default parameters followed by principle component analysis (PCA). The first 20 PCA were used for UMAP multi-dimensional reduction, based on the limited standard deviation in any further PCA. Graph based clustering with resolution of 1.0 was first used to identify clusters that also differentiated amongst sample types, before being annotated by grouping related clusters. Gene expression for ACE2, TMPRSS2, NRP1, and SARS-CoV-2 were assessed across samples as called by the Seurat function AverageExpression(). Comparison across samples and clusters was illustrated using VlnPlot() and DimPlot() functions.

### Shotgun proteomics of COVID-19 patient samples

The plasma of severe COVID-19 patients (N = 5) shown in Table 1 were collected and subjected to quantitative proteomics. Samples were lysed in 1% SDS, 100mM ammonium bicarbonate, 1mM EDTA, and 1X protease inhibitors. Samples were sonicated on ice, centrifuged at 10,000*g* and stored at-80°C until TMT-6plex™ Isobaric Labeling. Protein concentrations were determined by a NanoDrop 2000 spectrophotometer at 280nm. 200*μ*g of protein was mixed with lysis buffer (1% SDS, 100mM ammonium bicarbonate, 1mM EDTA, and 1X protease inhibitors) to a final volume of 100*μ*L. Samples were subjected to a quantitative proteomics workflow as per supplier (Thermo Fisher) recommendations. Samples were reduced in 200mM tris(2-carboxyethyl)phosphine (TCEP), for 1h at 55°C, reduced cysteines were alkylated by incubation with iodoacetamide solution (50mM) for 20min at room temperature. Samples were precipitated by acetone/methanol, and 600*μ*L ice-cold acetone was added followed by incubation at-20°C overnight. A protein pellet was obtained by centrifugation (8,000*g*, 10min, 4°C) followed by acetone drying (2min). Precipitated pellet was resuspended in100 *μ*L of 50mM triethylammonium bicarbonate (TEAB) buffer followed by tryptase digestion (5*μ*g trypsin per 100*μ*g of protein) overnight at 37°C. TMT-6plex™ Isobaric Labeling Reagents (90061, Thermo Fisher) were resuspended in anhydrous acetonitrile and added to each sample (41*μ*L TMT-6plex™ per 100*μ*L sample) and incubated at room temperature for 1h. The TMT labeling reaction was quenched by 2.5% hydroxylamine for 15min at room temperature. TMT labeled samples were combined and acidified in 100% trifluoroacetic acid to pH < 3.0 and subjected to C18 chromatography (Sep-Pak) according to manufacturer recommendations. Samples were stored at-80°C before lyophilization, followed by resuspension in 1% formic acid before liquid chromatography and tandem mass spectrometry analysis.

### Liquid Chromatography and Mass Spectrometry (LC-MS/MS)

Tryptic peptides were analyzed on an Orbitrap Fusion Lumos Tribrid mass spectrometer (Thermo Scientific) operated with Xcalibur (version 4.0.21.10) and coupled to a Thermo Scientific Easy-nLC (nanoflow liquid chromatography) 1200 System. Tryptic peptides (2*μ*g) were loaded onto a C18 trap (75*μ*m x 2cm; Acclaim PepMap 100, P/N 164946; ThermoFisher) at a flow rate of 2*μ*L/min of solvent A (0.1% formic acid in LC-MS grade H2O). Peptides were eluted using a 120min gradient from 5 to 40% (5% to 28% in 105min followed by an increase to 40% B in 15min) of solvent B (0.1% formic acid in 80% LC-MS grade acetonitrile) at a flow rate of 0.3*μ*L/min and separated on a C18 analytical column (75*μ*m x 50cm; PepMap RSLC C18; P/N ES803A; ThermoScientific). Peptides were then electrosprayed using 2.1kV voltage into the ion transfer tube (300°C) of the Orbitrap Lumos operating in positive mode. The Orbitrap first performed a full MS scan at a resolution of 120,000 FWHM to detect the precursor ion having a m/z between 375 and 1,575 and a +2 to +4 charge. The Orbitrap AGC (Auto Gain Control) and the maximum injection time were set at 4 × 10^5^ and 50ms, respectively. The Orbitrap was operated using the top speed mode with a 3 second cycle time for precursor selection. The most intense precursor ions presenting a peptidic isotopic profile and having an intensity threshold of at least 2 × 10^4^ were isolated using the quadrupole (Isolation window (m/z) of 0.7) and fragmented using HCD (38% collision energy) in the ion routing multipole. The fragment ions (MS2) were analyzed in the Orbitrap at a resolution of 15,000. The AGC and the maximum injection time were set at 1 × 10^5^ and 105ms, respectively. The first mass for the MS2 was set at 100 to acquire the TMT reporter ions. Dynamic exclusion was enabled for 45 seconds to avoid of the acquisition of same precursor ion having a similar m/z (plus or minus 10ppm).

## Data and code availability

The analysis scripts and reference genomes can be accessed at https://github.com/BiernaskieLab/scRNA-seq-SARS-CoV2-Viremia. All scRNAseq datasets generated in this study have been deposited in GEO under the following accession numbers.

To review GEO accession GSE151969 (Defining the global peripheral blood leukocyte response in severe COVID-19 patients admitted to the ICU with lung failure using single cell transcriptomics, 10 samples): Go to https://www.ncbi.nlm.nih.gov/geo/query/acc.cgi?acc=GSE151969 To be released, please contact the authors to request early access.

To review GEO accession GSE156639 (HUVEC response to SARS-CoV-2 and OC43 gene expression, 3 samples): Go to https://www.ncbi.nlm.nih.gov/geo/query/acc.cgi?acc=GSE156639 To be released, please contact the authors to request early access.

## Supplementary Materials

Table S1. Patient demographics

Table S2. Single cell RNAseq sample data

Table S3. Average gene expression per cell for each sample

Table S4. SARS-CoV-2 positivity across cell types

## Author Contributions

NR – conceived of the study, processed samples for 10X genomics, analysed scRNAseq, wrote the manuscript

AJ – pre-processed all scRNAseq data

CR, RPM, JAC – provided hCov-OC43, and conducted HUVEC transductions, and provided critical virology expertise

EL – processed samples for 10X genomics

SS – constructed all custom reference genomes LGA, AD – conducted proteomics and analysis

BY – enrolled patients, collected samples, cowrote the manuscript

JB – supervised experiments, cowrote the manuscript

