## Supplementary Tables for "SARS-CoV-2 infection of circulating immune cells is not responsible for virus dissemination in severe COVID-19 patients"

### Supplementary Information

Table 1. Patient demographics

| Sample | Sex | Age | PCR-based SARS-CoV-2 result |
|---|---|---|---|
| UC_4 | Female | 50 | Positive |
| UC_5 | Male | 52 | Positive |
| UC_6 | Male | 58 | Positive |
| UC_7 | Male | 67 | Positive |
| UC_8 | Male | 70 | Positive |

Table 2. Single cell RNAseq sample data

| Sample | Pre-normalization Total Reads/Cell (MT412228) | Number of cells pre-normalization and subsetting (MT412228) | Number of cells post-normalization and QC subsetting (MT412228) | Number of cells post-normalization and QC subsetting (MN908947.3) |
|---|---|---|---|---|
| UC_4 | 51,697 | 8,308 | 7,892 | 7,892 |
| UC_5 | 50,246 | 8,075 | 7,832 | 7,832 |
| UC_6 | 51,346 | 4,709 | 4,653 | 4,653 |
| UC_7 | 70,706 | 3,293 | 2,835 | 2,835 |
| UC_8 | 61,598 | 3,691 | 3,565 | 3,565 |
| BALF_46_CD45_HC | 80,975 | 11,200 | 9,626 | 9,626 |
| BALF_47_ CD45_HC | 86,350 | 866 | 519 | 519 |
| BALF_48_ CD45_HC | 82,126 | 6,474 | 2,643 | 2,643 |
| BALF_49_severe | 167,131 | 2,383 | 1,855 | 1,858 |
| BALF_50_ severe | 198,401 | 6,936 | 3,222 | 3,219 |
| BALF_51_ severe | 83,393 | 2,469 | 2,278 | 2,277 |
| BALF_54_mild | 225,422 | 8,308 | 2,690 | 2,684 |
| BALF_55_mild | 119,999 | 20,276 | 19,145 | 19,104 |
| BALF_56_ severe | 67,867 | 17,374 | 16,278 | 16,291 |
| BALF_57_mild | 323,279 | 10,601 | 8,567 | 8,581 |
| BALF_58_ severe | 72,607 | 4,158 | 3,497 | 3,495 |
| BALF_59_ severe | 304,255 | 4,495 | 3,151 | 3,152 |

Table 3. Average gene expression per cell for each sample.

| Sample | SARS-CoV-2 | ACE2 | TMPRSS2 | NRP1 |
|---|---|---|---|---|
| UC_4 | 0.00 | 0.00 | 0.00 | 0.00 |
| UC_5 | 0.00 | 0.00 | 0.00 | 0.00 |
| UC_6 | 0.00 | 0.00 | 0.00 | 0.00 |
| UC_7 | 0.00 | 0.00 | 0.00 | 0.00 |

| | | | | |
|---|---|---|---|---|
| UC_8 | 0.00 | 0.00 | 0.00 | 0.00 |
| BALF_46_CD45_HC | 0.00 | 0.00 | 0.00 | **0.33** |
| BALF_47_CD45_HC | 0.00 | 0.00 | 0.00 | **0.03** |
| BALF_48_CD45_HC | 0.00 | 0.00 | **0.01** | **0.74** |
| BALF_49_severe | 0.00 | **0.01** | **0.06** | **0.19** |
| BALF_50_severe | **0.42** | **0.01** | **0.05** | **0.22** |
| BALF_51_severe | **1.49** | **0.01** | **0.05** | **0.12** |
| BALF_54_mild | **2.50** | 0.00 | **0.01** | **0.04** |
| BALF_55_mild | **0.01** | 0.00 | **0.01** | **0.12** |
| BALF_56_severe | **0.42** | 0.00 | **0.03** | **0.39** |
| BALF_57_mild | 0.00 | 0.00 | **0.01** | **0.34** |
| BALF_58_severe | 0.00 | 0.00 | **0.02** | **0.14** |
| BALF_59_severe | 0.00 | 0.00 | **0.02** | **0.17** |

Table 4. SARS-CoV-2 positivity across cell types.

| **Cell type** | **CoVNeg** | **CoVPos** |
|---|---|---|
| Mac_1 | 34,315 | 755 |
| Mac_2 | 19,002 | 1 |
| UC_TCells | 15,111 | 0 |
| Neutrophils | 9,977 | 167 |
| TCells | 9,665 | 75 |
| UC_BCells | 1,875 | 0 |
| BCells | 2,076 | 55 |
| Epithelial | 2,414 | 148 |
| DC | 1,215 | 8 |
| NK | 2,044 | 12 |
| UC_Monocytes | 863 | 0 |
| UC_Platelets | 437 | 0 |
| Myeloid | 36 | 1 |
